# Fast layer-fMRI VASO with short stimuli and event-related designs at 7T

**DOI:** 10.1101/2023.03.15.532735

**Authors:** Sebastian Dresbach, Renzo Huber, Omer Faruk Gulban, Rainer Goebel

## Abstract

Layers and columns are the dominant processing units in the human (neo)cortex at the meso-scopic scale. While the blood oxygenation dependent (BOLD) signal has a high detection sensitivity, it is biased towards unwanted signals from large draining veins at the cortical sur-face. The additional fMRI contrast of vascular space occupancy (VASO) has the potential to augment the neuroscientific interpretability of layer-fMRI results by means of capturing com-plementary information of locally specific changes in cerebral blood volume (CBV). Specifically, VASO is not subject to unwanted sensitivity amplifications of large draining veins. Because of constrained sampling efficiency, it has been mainly applied in combination with efficient block task designs and long trial durations. However, to study cognitive processes in neuroscientific contexts, or probe vascular reactivity, short stimulation periods are often necessary. Here, we developed a VASO acquisition procedure with a short acquisition period (895 ms volume acqui-sition) and sub-millimetre resolution. During visual event-related stimulation, we show reliable responses in visual cortices within a reasonable number of trials (∼20). Furthermore, the short TR and high spatial specificity of our VASO implementation enabled us to show differences in laminar reactivity and onset times. Finally, we explore the generalizability to a different stimulus modality (somatosensation). With this, we showed that CBV-sensitive VASO provides the means to capture layer-specific haemodynamic responses with high spatio-temporal resolution and is able to be used with event-related paradigms.

## Introduction

Cortical layers give rise to fundamental processing units like the cortical microcircuit (Bastos et al., 2012; Douglas and Martin, 2004) and may inform about feedforward and feedback characteristics of brain regions in human and animal brains (Felleman and Van Essen, 1991; Markov et al., 2014). Due to technological advancements in the last decade, neuroscientists are now able to study these structures non-invasively in humans using functional magnetic resonance imaging (fMRI) at high resolutions (<1*μ*L voxel volume, Polimeni et al., 2010). However, when approaching this spatial scale, several challenges remain. Most notably, the so-called “draining vein” effect is known to have a major impact on the fMRI signals measured across cortical depths when using the popular gradient echo (GE) blood-oxygenation level dependent (BOLD) sequences (Turner, 2002). Specifically, the venous blood is drained from deep to superficial layers within the cortex leading to a spatial displacement of neuronal responses measured indirectly by the BOLD signal. This decreases the effective spatial specificity, despite having small voxel sizes (De Martino et al., 2013; Menon and Goodyear, 1999; Self et al., 2019).

To counter the neurovascular confounds in the BOLD signal, additional measurements of cerebral blood flow (CBF) and cerebral blood volume (CBV) have been proposed. These approaches promise higher specificity to the neural site of activation by being less affected by the draining veins (L. Huber et al., 2019). One of the most widely used non-invasive measurements of CBV is vascular space occupancy (VASO, Hua et al., 2013; Lu et al., 2003). Slice-selective slab-inversion (SS-SI) VASO (L. Huber et al., 2015, henceforth referred to as VASO) has been used to study fundamental neuroscientific mechanisms on the mesoscopic scale (L. Huber et al., 2017; Persichetti et al., 2020; Y. Yu et al., 2019). However, the challenging contrast-to-noise-ratio of sub-millimeter fMRI and the constrained sampling efficiency of VASO have so far hindered wide adaptation of its application in neuroscientific application studies with fast stimulus designs. While previous studies suggest that slow event-related paradigms with 6 s long task durations are feasible with layer-fMRI VASO (Persichetti et al., 2020), it is not clear if VASO can be pushed to even faster designs. Furthermore, it has not been clear until now if and how much the detection sensitivity is compromised in layer-fMRI VASO for event-related design tasks compared to commonly used block-design tasks.

Event-related task designs are widely-adopted in fMRI to conduct cognitive neuroscience research (Huettel, 2012). This is because event-related task designs provide flexibility in stim-ulus presentation to control for a wide range of confounds like psychological effects (carryover effects, habituation, anticipation, etc., Rosen et al., 1998). Other uses of event-related task designs revolve around studying the individual trials e.g. to characterize learning, correlating responses to behavioral variables (e.g. reaction time), or post-hoc sorting of trials based on e.g. subjective perceptions [e.g. Formisano et al., 2002; Heynckes et al., 2023; Schneider et al., 2019). Although some of these aspects are also achievable with longer stimulus durations (e.g. 6 second long randomized stimuli as implemented by Persichetti et al. (2020), most other benefits of event-related task designs can only be exploited by using shorter stimulus durations.

In this study, we implemented, tested, and validated a VASO sequence protocol with a short repetition time (TR, 2.57 s) that provides the means to acquire layer-specific haemodynamic responses with high temporal resolution and sufficient coverage to capture multiple distant brain areas. For this, we used high resolution (0.88 mm3 isotropic nominal voxel size) fMRI at ultra-high field strength (7 Tesla). We further characterized the sub-millimeter CBV responses and their detection sensitivity to block and event-wise stimulation. Third, we explored the generalizability to different stimulus modalities (vision and somatosensation). As the VASO sequence yields VASO and BOLD images, we made use of both contrasts to present our results, which offers complementary information. Using our new sequence protocol, we showed that it is possible to capture CBV-VASO responses to stimuli with short durations in a fast event-related design within a reasonable amount of trials. Our results form the foundation of a new avenue for further adoption of VASO in cognitive neuroscience and physiology studies.

## Materials and Methods

### Participants

10 right handed participants (2 female, age range: 23-31 years, mean age: 27.9 years), with no neurological damage participated in the study after providing written informed consent. The paradigm was approved by the local Ethics Review Committee for Psychology and Neuroscience (ERCPN) at Maastricht University (ERCPN refnr 180 03 06 2017).

### Imaging parameters

All participants underwent scanning using a ”classical” 7T Magnetom whole body scanner (Siemens Healthineers, Erlangen, Germany), equipped with a 1Tx, 32Rx head RF coil (Nova Medical, Wilmington, MA, USA) at Scannexus B.V. (Maastricht, The Netherlands). Functional scans were performed with 3D echo-planar imaging (Poser et al., 2010) VASO (L. Huber et al., 2014) and a nominal resolution of 0.875 × 0.875 × 0.88 mm3, 16 slices, TI/volumeTR/pairTR/TE = 1272/895/2570/18 ms, partial Fourier factor = 6/8 using projection onto convex sets (POCS, Nakamura et al., 2016), reconstruction with 8 iterations, FLASH GRAPPA 3 (Talagala et al., 2016), varying flip angles between 26 and 90°, read bandwidth = 1266 Hz/Px, first phase encoding direction along anterior-posterior axis with axial-like tilted slice orientation, matrix size = 152 × 148, field of view (FOV) = 133 × 129.542 mm in read and first phase encoding directions respectively. We used the vendor-provided ‘improved GRAPPA WS’ algorithm with at least 1000 fold over regularization and small GRAPPA kernels of 2×3 (phase × read). Parameters are summarized in **Figure 1B**, a scan protocol is available on github (https://github.com/sdres/sequences/blob/master/20211022 Seb.pdf) and the sequence is avail-able for VB17B via SIEMENS’ C2P app store Teamplay.

**Figure 1:**
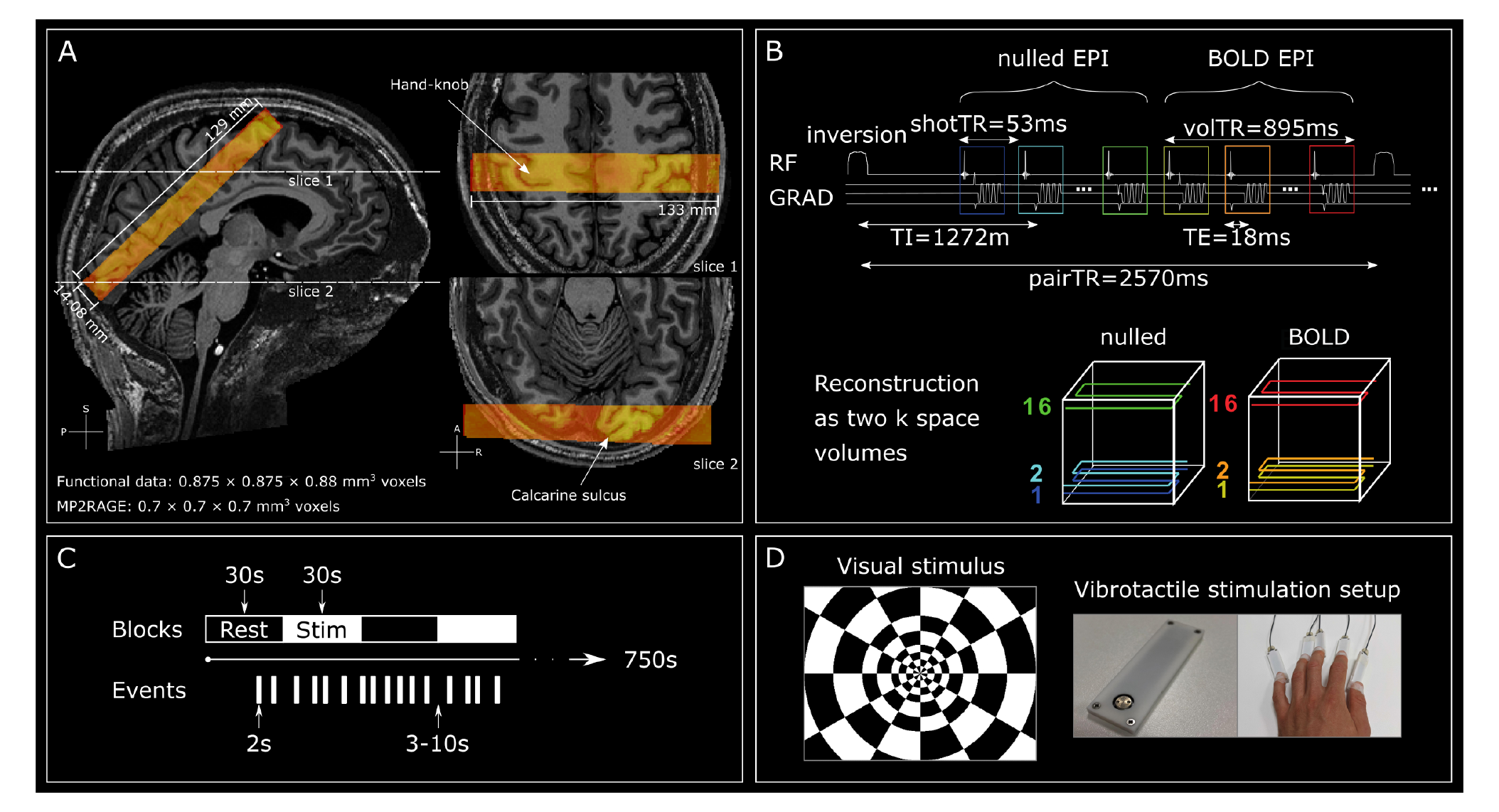
fMRI data acquisition and stimulation protocol. **A** Functional EPI data (warm colors) overlaid on MP2RAGE UNI image (sub-07). The fMRI slab covered bilateral calcarine sulci (lower right) and when possible the hand area of the right postcentral gyrus (upper right). **B** Functional EPI data acquisition details giving a schematic overview of relevant imaging parameters. (Note: nulled and BOLD images are acquired in an interleaved fashion. Both volumes are later used to derive the ‘VASO’ images.) **C** Illustration of block and event-wise stimulation runs. Block-wise stimulation consisted of 30s on-off periods. Event-wise stimuli were 2 seconds in duration separated by inter-trial intervals of 3-10 seconds which were randomly chosen from a uniform distribution. **D** Means of stimulation. Left: Flickering checkerboard. Contrast reversals were displayed at 16 Hz. Right: A single piezoelectric stimulation device. The silver disc vibrates at 25 Hz and stimulates the fingertips. On the right, 5 devices are attached to all fingers of the left hand with elastic tape.

Two participants were invited for a second session, in which we compared the above protocol to a version of the sequence with a longer TR. Specifically, we inserted 500 ms delays between the image readouts with and without blood-nulling, as well as between the readout of the BOLD (also referred to as ‘not-nulled’) image and the consecutive inversion pulse. Thus, the pair TR was 3570 ms, while all other parameters remained equal.

Slice position and orientation were chosen individually for each participant. We covered bilateral calcarine sulci and the hand area of the right postcentral gyrus (both indicated in **Figure 1A**). We localized the area of interest in the right somatosensory cortex based on the position of the ‘hand-knob’ area on the precentral gyrus, opposite of which the finger representations are located on the postcentral sulcus. Note that, targeting both the visual and the somatosensory cortex turned out to be challenging without significant fold-over artifacts due to the small FOV and limited number of slices. In cases where coverage of both visual and somatosensory cortices was not possible (n = 5), we prioritized the visual cortex.

Finally, we either acquired high-resolution whole-brain anatomical images (0.7 mm isotropic, 240 slices, TI1/TI2/TR/TE = 900/2750/5000/2.47 ms, FA1/FA2 = 5°/3°, bandwidth = 250 Hz/Px, acceleration factor = 3, FoV = 224 × 224 mm) using MP2RAGE (Marques et al., 2010), or images were available from previous scans with similar parameters. For one participant, we neither acquired MP2RAGE data due to time constraints, nor was there previous data available.

### Stimuli

Stimuli consisted of flickering checkerboards presented at 16 Hz and vibrotactile stimulation of all 5 digit-tips (left hand) by means of a piezoelectric stimulator at 25 Hz (mini PTS system, Dancer Design, UK). Both means of stimulation are displayed in **Figure 1D**. Stimuli were presented either in blocks or as events while the run duration was kept to 12.5 minutes in both cases (**Figure 1C**). We acquired 1-2 blocked-and 2-4 event wise stimulation runs per participant for the main experiment. For the comparison between long and short TR protocols, we acquired 4 runs of block-wise stimulation runs (2 per TR duration). During block-wise stimulation runs, we presented stimuli for 30 s with 30 s of resting fixation in between blocks, resulting in 12 blocks per run with an additional rest period before the first stimulation block. Event-wise stimulation runs started with a 19 s baseline period and then stimuli were presented for 2 s per trial with inter-trial intervals between 3 and 10 s chosen from a uniform distribution, resulting in 84 trials per run (12.5 minutes). These highly irregular inter-trial intervals were chosen to accurately capture the haemodynamic responses of BOLD and VASO. The stimulation pattern was generated once and then used for all runs in order to allow averaging. The stimulation was controlled via Psychopy 3 v2020.2.4 (J. Peirce et al., 2019; J. W. Peirce, 2007) scripts, which can be found at: <https://github.com/sdres/eventRelatedVASO/tree/main/code/stimulation>.

In the scenario described above, visual and tactile stimulation was always presented concurrently. For five participants, we deviated from this stimulation pattern. During one session of one participant, the vibrotactile stimulation device was not available. Therefore, we only presented visual stimuli during that session. For four other participants, we randomized visual only and visuo-tactile stimulation during the event-related runs. This way, we aimed to disentangle potential multisensory and physiological effects of laminar reactivity.

### Processing

The analysis code is available at <https://github.com/sdres/eventRelatedVASO>. Briefly, initial motion-correction was performed within runs for nulled and not-nulled time-series separately using ANTsPY v0.2.7 (Avants et al., 2011). We used an automatically generated brain mask (3dAutomask in AFNI 22.1.13, Cox, 1996) to calculate cost functions in brain tissue only. We then computed T1w images in EPI-space derived from the combined nulled and BOLD im-ages for each run (3dTstat with -cvarinv) using AFNI, performed bias field correction using ANTS v2.3.5.dev238-g1759e (N4BiasFieldCorrection, Tustison et al., 2010) and registered the resulting image to the first event-wise stimulation run using ANTsPY. We then performed the motion correction again from scratch, while concatenating the transformation matrices from within and between run registration. Due to the small imaging slab, registering different ses-sions of the same participant was challenging. Therefore, we treated them as independent datasets. However, as they were only acquired to assess differences between long and short TR flavors of the protocol, this does not influence the results of our main experiment.

The VASO sequence acquires images with BOLD and CBV weighting concomitantly in an interleaved fashion. Therefore, we temporally upsampled the motion corrected data with a factor of 2 (3dUpsample in AFNI with 7th order polynomial interpolation). We then per-formed BOLD correction and computed standard measures for quality control (tSNR, skew, and kurtosis) using LayNii (v2.2.1, LN BOCO and LN SKEW respectively; L. (Huber et al., 2021). General linear model (GLM) analyses were performed in FSL (Woolrich et al., 2001, v6.0.5.2, contrast: stimulation vs rest). MP2RAGE UNI images were registered to the T1w EPI image of the first event-wise stimulation run, first, using manual alignment in ITK-SNAP (v3.8.0, Yushkevich et al., 2006, then an automatic alignment (affine transformation, mutual information cost metric) while using the brain mask generated for the motion correction to guide registration. Finally, anatomical images were registered nonlinearly to the T1 weighted image in EPI space using ANTS.

Borders between gray matter (GM) and cerebrospinal fluid (CSF) and between GM and white matter (WM) of ROIs were manually drawn in FSLeyes v0.31.2 (McCarthy, 2023) based on the activation in response to block-wise stimulation and the registered MP2RAGE UNI contrast. For the participant, where no MP2RAGE data was available, we drew the borders based on residual contrast of the T1-weighted images directly in EPI space. All manually drawn masks are available alongside the raw data. Finally, layering was performed on spatially upsampled data (3dresample factor 5 in-plane) using the equidistant approach as implemented in LayNii (LN2 LAYERS). For the estimation of cortical depth profiles of GLM analyses, we defined 11 depth bins, while for the finite impulse response (FIR) analysis across cortical depth, we estimated three depth bins (deep, middle, superficial) to allow for appropriate averaging.

For the analysis of the block-wise stimulation runs, we ran a GLM with one predictor for the stimulation times, convolved with a standard gamma haemodynamic response function without temporal derivative (mean lag: 6 s, std. dev: 3 s). Here, we applied a high-pass filter (cutoff = 0.01 Hz) and no additional smoothing. To estimate the temporal response for blocked stimulation, we averaged epochs with stimulation with 4 volumes before stimulus onset and 8 volumes after cessation. The percent signal change was computed with respect to the 30 s rest-periods in between stimulation blocks.

Responses to event-wise stimulation runs were first estimated with a GLM, similar to the blocked stimulation for better comparison. We then ran a GLM analysis using finite impulse response models on the event-wise stimulation data. Here, we modeled 10 timepoints after stimulus onset, resulting in a window of 13.08 seconds.

To estimate the number of trials necessary to obtain a reliable signal, we extracted time-courses from the ROIs, epoched the data with a window from stimulus onset until 8 s after cessation. We then computed the overall epoch activity across all participants. For each run, we then computed a cumulative average for 1 up to 84 trials, where 84 was the number of trials in a run. For each number of averaged trials, we computed the sum of squares between the values for each timepoint separately and the average response across all participants. Based on anatomical landmarks, we will refer to ROIs in the visual cortex as V1 and ROIs on the postcentral gyrus as S1.

## Results

In the following, we will focus on the results in visual cortices as this was the primary area of interest of this study. The exploratory results for the somatosensory cortex will be described briefly afterwards. Finally, one participant (sub-10) consistently showed excessive head motion well above the voxel size and was therefore excluded from further analyses (**Supplementary Figure S1**).

### Block-wise stimulation

Regarding the main experiment, we first examined the results from block-wise stimulation for two reasons. First, we compared our results to previous research to see whether the results obtained with our fast protocol match the expected patterns. Second, we used these data to determine ROIs for further analyses.

As expected, block-wise stimulation elicited strong signal changes in the visual cortices for both BOLD and VASO in all participants. **Figure 2A** shows GLM activation in response to stimulation (z-scores, contrast: stimulation vs. rest) overlayed on an MP2RAGE UNI image for one individual. As expected, BOLD activation scores are generally higher than for VASO. On the other hand, the active voxels are mostly constrained to cortical GM for VASO, while BOLD shows largest z-scores in CSF. Based on this, we manually delineated ROIs from which we extracted signal changes over time and z-scores across cortical depth. As expected, the temporal profiles of the BOLD and VASO responses show sustained activity, peaking around 10s with an additional off-response after stimulus cessation (**Figure 2B**). Interestingly, we also observed a strong post-stimulus undershoot for BOLD, while this was less pronounced in VASO. Finally, **Figure 2C** shows the activation profile across cortical depths. Here, we see the well reproduced result of increasing activity towards the surface for BOLD and the inverted U-shaped activity pattern for VASO. These likely reflect the drainage of deoxygenated blood towards the pial surface and the thalamocortical input to layer 4, respectively (Felleman and Van Essen, 1991; Turner, 2002). Taken together, these results assured us that our protocol gives robust results with paradigms.

**Figure 2:**
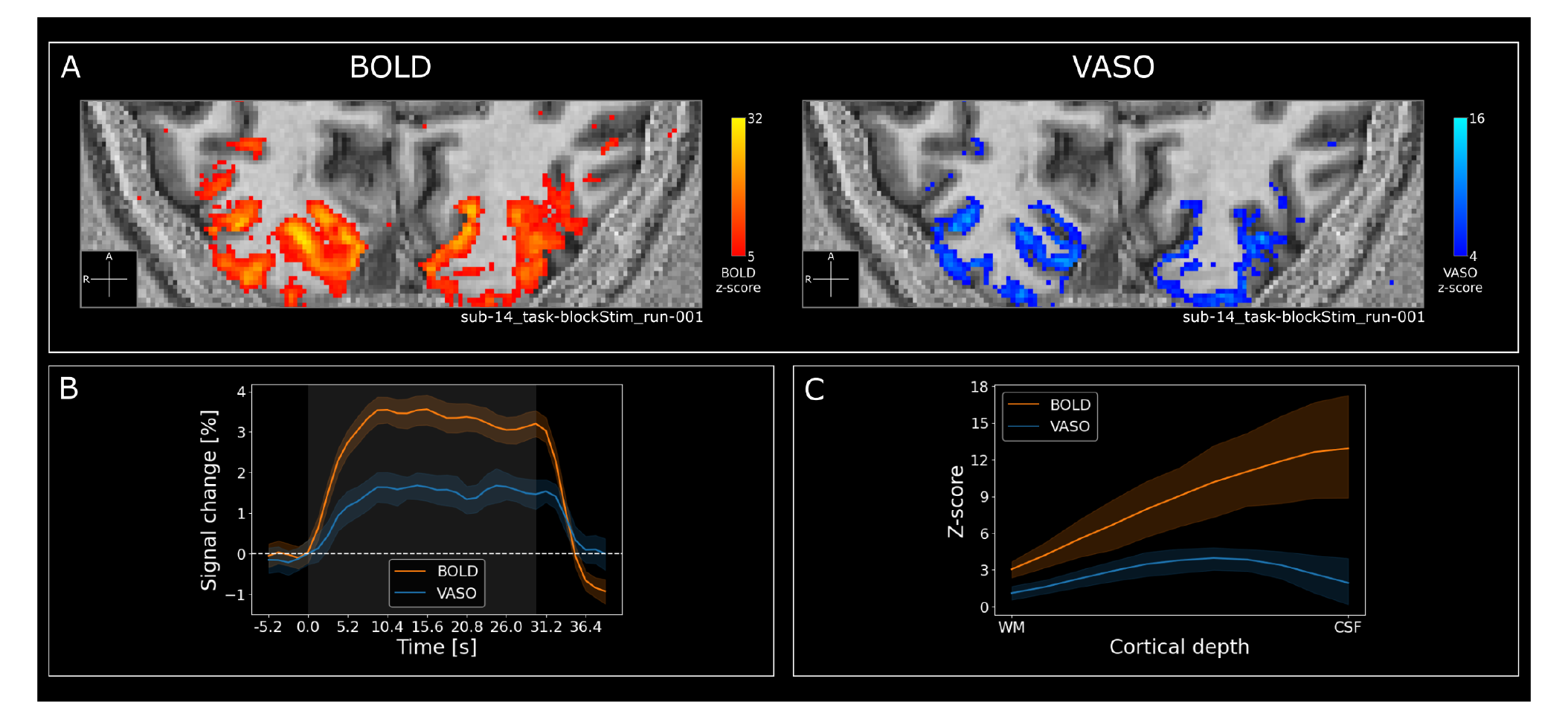
Block-wise stimulation results show robust activation. **A** GLM activation in response to stimulation (z-scores, contrast: stimulation vs. rest) overlayed on MP2RAGE UNI image. Note that here we show results from a single run (12.5 minutes). **B** Group-level averages showing the BOLD and VASO signal change across all layers in response to stimuli with a duration of 30s (shaded gray area). Colored shaded areas represent 95% confidence intervals across blocks. **C** Group-level layer profiles showing GLM activation in response to stimulation (z-scores, contrast: stimulation vs. rest) for BOLD and VASO in V1 across cortical depths. Colored shaded areas represent 95% confidence intervals across participants.

### Event-wise stimulation

Event-wise stimulation also elicited signal changes in the visual cortices for both BOLD and VASO in all participants, albeit to a lesser degree. **Figure 3A** shows GLM activation in response to stimulation (z-scores, contrast: stimulation vs. rest) for two runs of one individual overlayed on an MP2RAGE UNI image. Despite the lower z-scores compared to block-wise stimulation in both BOLD and VASO, clear patterns can be seen without the need for additional processing (e.g. denoising/ smoothing). Specifically, activated VASO voxels follow the cortical ribbon as expected, whereas BOLD responses are strongest at the cortical surface and in CSF.

**Figure 3:**
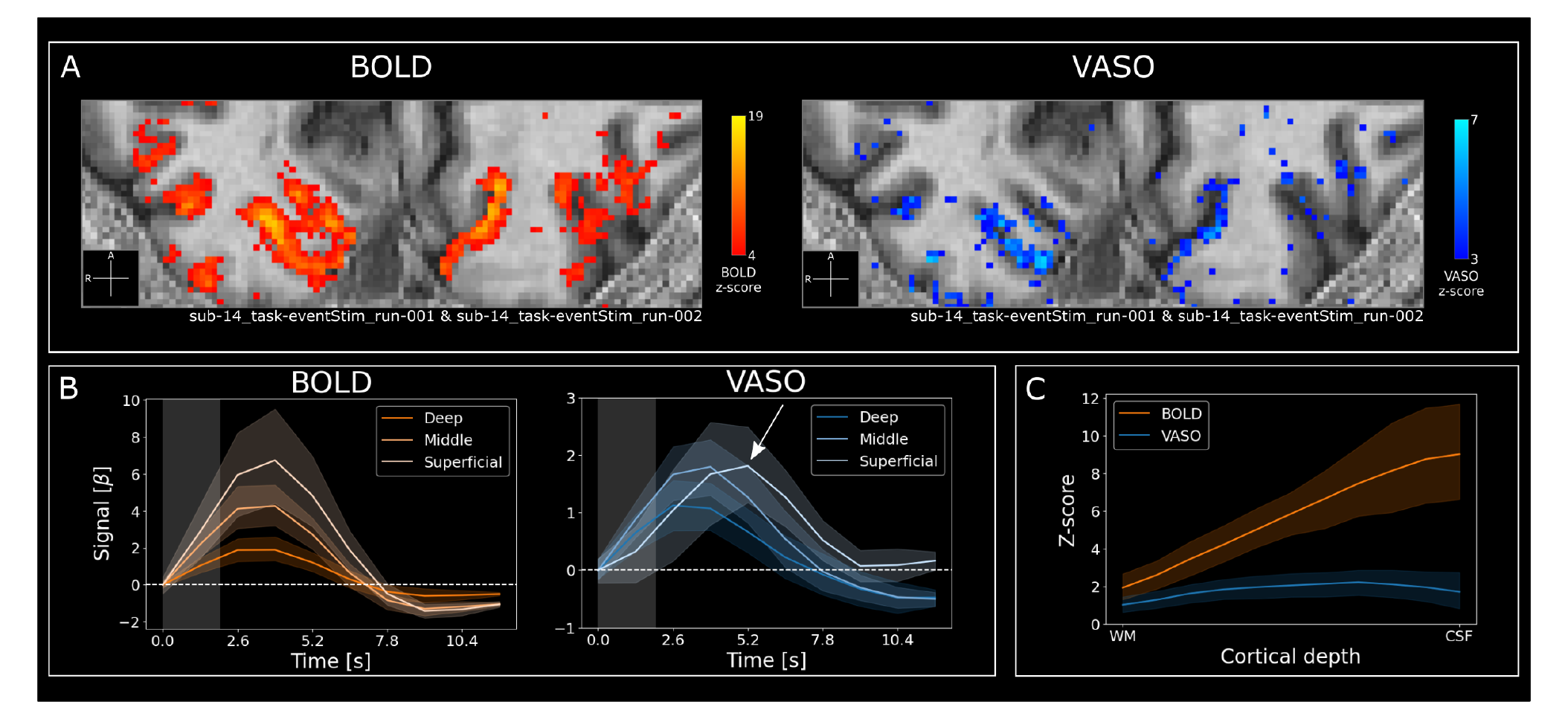
Event-wise stimulation results also show robust activation. **A** GLM acti-vation in response to event-wise stimulation (z-scores, contrast: stimulation vs. rest) in V1 of a representative participant (sub-14) overlayed on an MP2RAGE UNI image. Note that here we show data from two runs (25 minutes). **B** Group-level average responses showing the BOLD (left) and VASO (right) model fit (FIR model with 10 predictors) for three layers (deep, middle, superficial) in response to stimuli with a duration of 2 seconds (shaded gray area). The white arrow indicates the delayed peak for the superficial cortical depth in VASO. The colored shaded areas represent 95% confidence intervals across runs. **Supplementary Figure S2** shows the same plot for 3 individuals. **C** Group-level layer profiles showing GLM activation in response to stimulation of 2 seconds (similar to **Figure 3A**, z-scores, contrast: stimulation vs. rest) for BOLD and VASO in V1 across cortical depths. Colored shaded areas represent 95% confidence intervals across participants.

To investigate the haemodynamic response over time, we performed a deconvolution analysis using a FIR model as implemented in FSL with three layer bins (**Figure 3B**). Responses for both BOLD and VASO followed the expected haemodynamic response. As expected, BOLD signal changes increased from deep to superficial layers. This was less apparent in VASO. To our surprise, group-level VASO responses showed comparable signal changes for middle and superficial layers as opposed to the strongest response in middle layers for block-wise stimulation. This was highly variable across participants (**Supplementary Figure S2**). Most notably however, responses in superficial layers were delayed with respect to deep and middle layers (white arrow in the right panel of **Figure 3B**). This finding was highly consistent across participants, irrespective of superficial layer response amplitude (**Supplementary Figure S2**). Also note that the response in deep layers does not differ strongly between BOLD and VASO. This might indicate similar detection sensitivity between BOLD and VASO for GM close to the WM/GM border.

The stimulus consisted of visual and tactile stimulation and so, we tested whether the lam-inar timing differences might be explained by effects of multisensory integration. For this, we randomly presented visual or visuo-tactile stimuli in an event-related fashion in 4 participants. The results are shown in **Supplementary Figure S3**. Briefly, visual and visuo-tactile stimu-lation evoked very similar responses in superficial and middle layers for BOLD and VASO, thus ruling out multisensory effects as an explanation for the delayed peak in superficial layers for VASO. In deep layers, visuo-tactile stimulation showed a slightly prolonged response compared to visual stimulation only. This effect was present in the BOLD and VASO data, while being more pronounced in the latter. However, this effect is rather small (within error bars) and has to be interpreted with caution.

Finally **Figure 3C** shows the activation profile across cortical depths. Also here, the increasing activity towards the surface for BOLD is clearly visible. The inverted U-shaped activity pattern for VASO is also present albeit less clear than for the block-wise stimulation results. Taken together, these results show that it is possible to capture event-related responses to short stimuli using CBV-based VASO measurements.

### How many trials do we need?

**Figure 4A** shows the response stabilization when cumulatively averaging trials. Here, the average of a given number of trials is compared to the response across all participants. The number of average trials ranges from 1 to 84 (a run had a maximum of 84 trials). Thus, the last datapoint on the x-axis constitutes the difference between the average response of a run and the overall mean. While the difference between a few trials and the response across all participants is large, the response quickly approaches the overall average within a few more trials. Across participants, the 5% of the error dynamic range for VASO is reached when averaging 20 trials. The 5% of the error dynamic range for BOLD is reached when averaging 23 trials. The slightly higher number of averages needed for BOLD is due to the smaller dynamic range. Still, this shows that the noise level of our data is acceptable to efficiently capture the response to event-related stimuli without the need to average over long periods of time. Note that we used strong stimuli (flickering checkerboards) and did not perform any experimental manipulation. Most neuroscientific investigations use weaker tasks with control conditions that are very similar to the experimental condition. Therefore, in these experiments the number of presented stimuli will most likely have to be higher in order to reliably separate responses to the conditions. In **Figure 4B** we directly compared GLM activation in response to block- and event-wise stimulation for BOLD and VASO separately. In our BOLD data, we see lower signal intensity for event-than for block-wise stimulation. This decrease in activation scores is similar across cortical depths. In VASO, the pattern is more complex. While overall signal intensity does not differ much between block- and event-wise stimulation for deep and superficial layers, the group-level response peak is more biased towards the surface for event-wise stimulation. As discussed before, this response peak shows large variability across participants (see previous section and **Supplementary Figure S2**). Further, we aimed to estimate the change in detection sensitivity when employing block- vs. event-wise stimulation designs. From the data plotted in **Supplementary Figure 4B**, we approximated the area under the curve for block- and event-wise stimulation separately for BOLD and VASO, by summing the z-scores of all the layer-bins for block- and event-wise stimulation. We then subtracted the sum of the event-wise from the block-wise results and normalized them to the latter. Across subjects, detection sensitivity decreased 36 % in BOLD and 22 % in VASO. Note, we performed 1-2 block-wise and 2-4 event-wise stimulation runs per participant. Still, the direct comparison of block- and event-wise stimulation shows that it is possible to use event-related designs within an acceptable acquisition time of 25 minutes (see **Figure 4A**).

**Figure 4:**
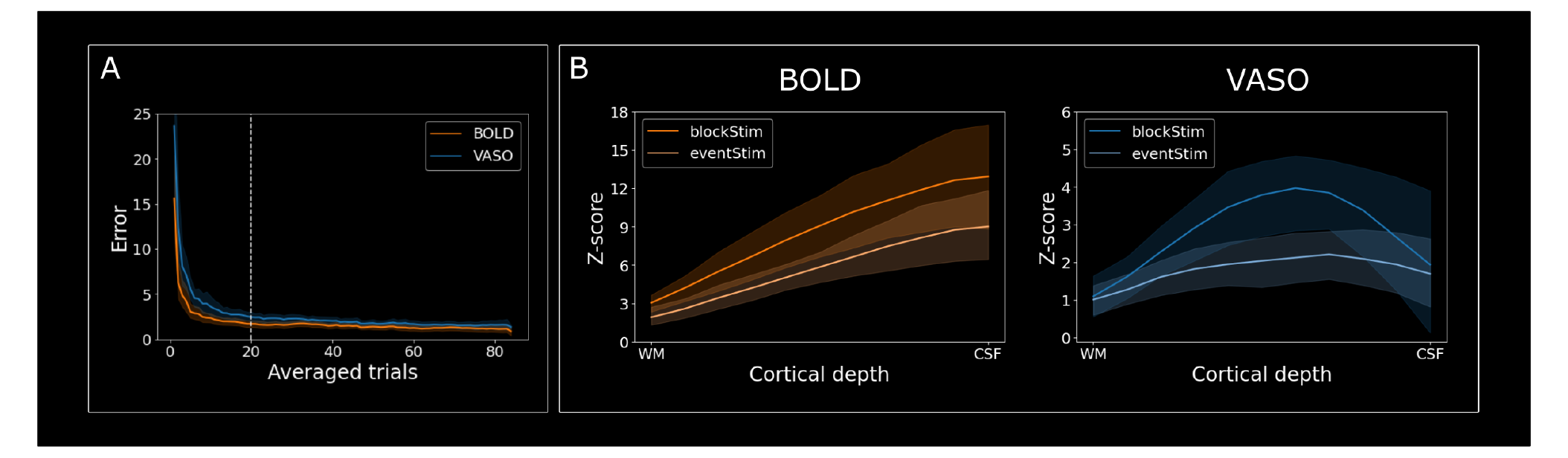
VASO responses stabilize after 20 averaged trials and event-wise stimulation yields lower but plausible activation compared to block-wise stimulation. **A** Response stabilization curves for BOLD and VASO. Note, a run had a maximum of 84 trials. Thus, the last datapoint on the x-axis constitutes the difference between the average response of a run and the overall mean. Dashed vertical line shows the 5% of the error dynamic range for VASO. **B** Data from **Figure 2C** and **Figure 3C** are replotted to compare block- with event-wise stimulation directly for BOLD (left) and VASO (right) separately. Note, we performed 1-2 block-wise and 2-4 event-wise stimulation runs per participant.

### Exploratory results in the somatosensory cortex

In 5 sessions we succeeded in including both the visual and right somatosensory cortex in the fMRI imaging slab. Therefore, we can explore the applicability of event-related stimulation using VASO in S1. Here, we obtained similar results as in V1, albeit less clear.

In general, fingertips are represented sparsely along the cortical sheet. Therefore, even if we managed to include S1 in some participants, we did not necessarily include all five finger representations for each of them. Therefore, we limited our analysis to the likely site of one fingertip. **Figure 5A** shows GLM activation in S1 in response to block-wise stimulation (z- scores, contrast: stimulation vs. rest) overlayed on an MP2RAGE UNI image. For the BOLD data, we can see clear activation. For VASO, only a few voxels exceed the detection threshold. Nevertheless, we were able to observe sustained periods of elevated signal in both imaging modalities **Figure 5B**) and group-level layer profiles showing GLM activation in response to block-wise stimulation (**Figure 5C**). Also the event-wise stimulation results show similar qualities of the responses in V1, both with higher noise levels (**Figure 5D&E**). Namely, just like in V1, in S1 VASO responses in superficial layers show indications of a later peak compared to deeper layers. These results point towards the applicability of event-wise stimulation using VASO in S1.

**Figure 5:**
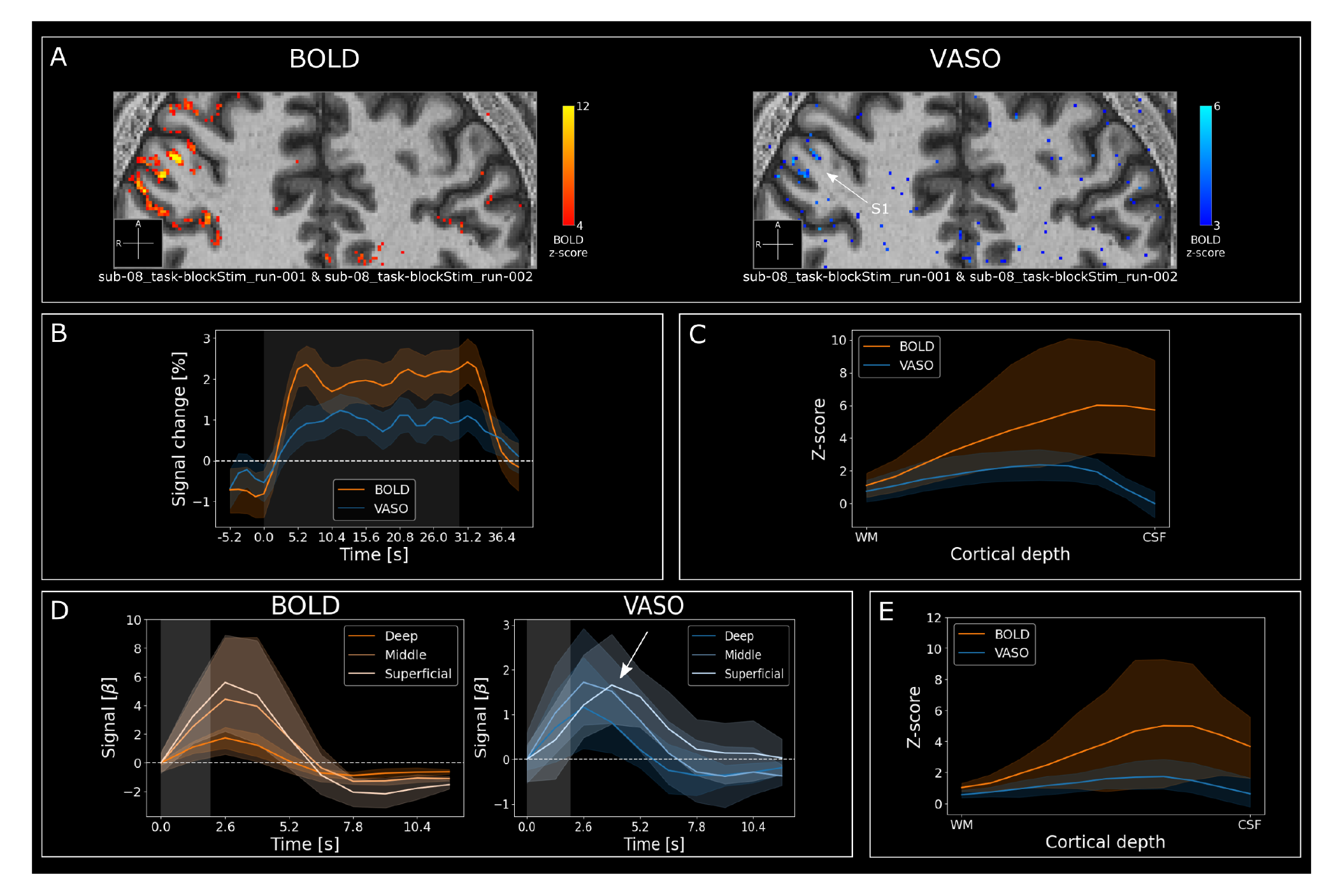
Exploratory results point towards feasibility of event-related designs in S1 using VASO. **A** GLM activation in response to block-wise stimulation (z-scores, contrast: stimulation vs. rest) overlayed on MP2RAGE UNI image. Note that here we show data from two runs (25 minutes). **B** Group-level (n = 5) averages showing the BOLD and VASO signal change across all layers in response to stimuli with a duration of 30s (shaded gray area). Colored shaded areas represent 95% confidence intervals across blocks. **C** Group-level (n = 5) layer profiles showing GLM activation in response to block-wise stimulation (z-scores, contrast: stimulation vs. rest) for BOLD and VASO in S1 across cortical depths. Colored shaded areas show 95% confidence intervals across participants. **D** Group-level (n = 5) average responses showing the BOLD (left) and VASO (right) model fit (FIR model with 10 predictors) for three layers (deep, middle, superficial) in response to stimuli with a duration of 2 seconds (shaded gray area). The colored shaded areas represent 95% confidence intervals across runs. **E** Group-level (n = 5) layer profiles showing GLM activation in response to stimulation of 2 seconds (similar to **Figure 3C**, z-scores, contrast: stimulation vs. rest) for BOLD and VASO in S1 across cortical depths. Colored shaded areas represent 95% confidence intervals across participants.

### Comparing short and long TR protocols

Because we stepped into uncharted territory with our short acquisition times, we wanted to test whether the activation profiles across cortical depth are influenced by the short TR compared to conventional acquisitions with longer intervals between readouts. SS-SI VASO is based on the assumption that the intravascular water magnetization in the imaging slab has been inverted only once (L. Huber et al., 2014). This means that blood should be refilled in the period between two consecutive inversion pulses (here 2.57s). While this condition is expected to be fulfilled for the thin imaging slab used here, this has not been validated for the unprecedented fast imaging of this study. Therefore, we directly compared data from our new protocol and a more conservative approach. In the latter, we added 500ms delays between the nulled and BOLD acquisitions and between the BOLD and the consecutive inversion pulse. As this decreased the efficiency of our temporal sampling, we tested this approach using block-wise stimulation (30s on-off) with our visuo-tactile paradigm. We found several differences between the two types of acquisitions. First, we observed a reduction of T1w contrast for short compared to long TR acquisitions (**Figure 6A**). While we can clearly see the borders between GM and CSF/WM, for long TRs, the borders between GM and WM are less visible for our short TR protocol. This different structural T1-contrast across TRs is expected due to the different z- magnetization steady-state in absence of a relaxation delay before each inversion pulse. Because of the decreased contrast in short TR protocols, we used the whole brain anatomical MP2RAGE images and registered them to the functional data in order to delineate GM/WM and GM/CSF borders as commonly done in GE-BOLD type acquisitions. Second, the tSNR of the BOLD time-course was slightly higher for the long TR, while the VASO tSNR was largely unaffected (Figure 6B). The higher tSNR for BOLD for long TRs is likely due to the further signal relaxation of the longitudinal tissue magnetization subsequent to the VASO readout. Finally, the statistical GLM activation in response to stimulation (z-scores, contrast: stimulation vs. rest) was slightly higher for long- than short TRs (**Figure 6C**). However, this difference was rather uniform across cortical depths. The slightly lower z-scores are to be expected due to the lower tSNR in BOLD. Evidently, this hampered the fit of the GLM for the BOLD data. The differences we see for the VASO data can be attributed to the dynamic division of BOLD and nulled data during the BOLD-correction step of the VASO data. Here, a higher noise level of the divisor will also enhance noise in the resulting signal. In conclusion, as can be seen in **Figure 6C**, the layer-profiles for long and short TRs were qualitatively not substantially different thus ensuring comparable signal origin and applicability of the VASO assumptions (L. Huber et al., 2014).

**Figure 6:**
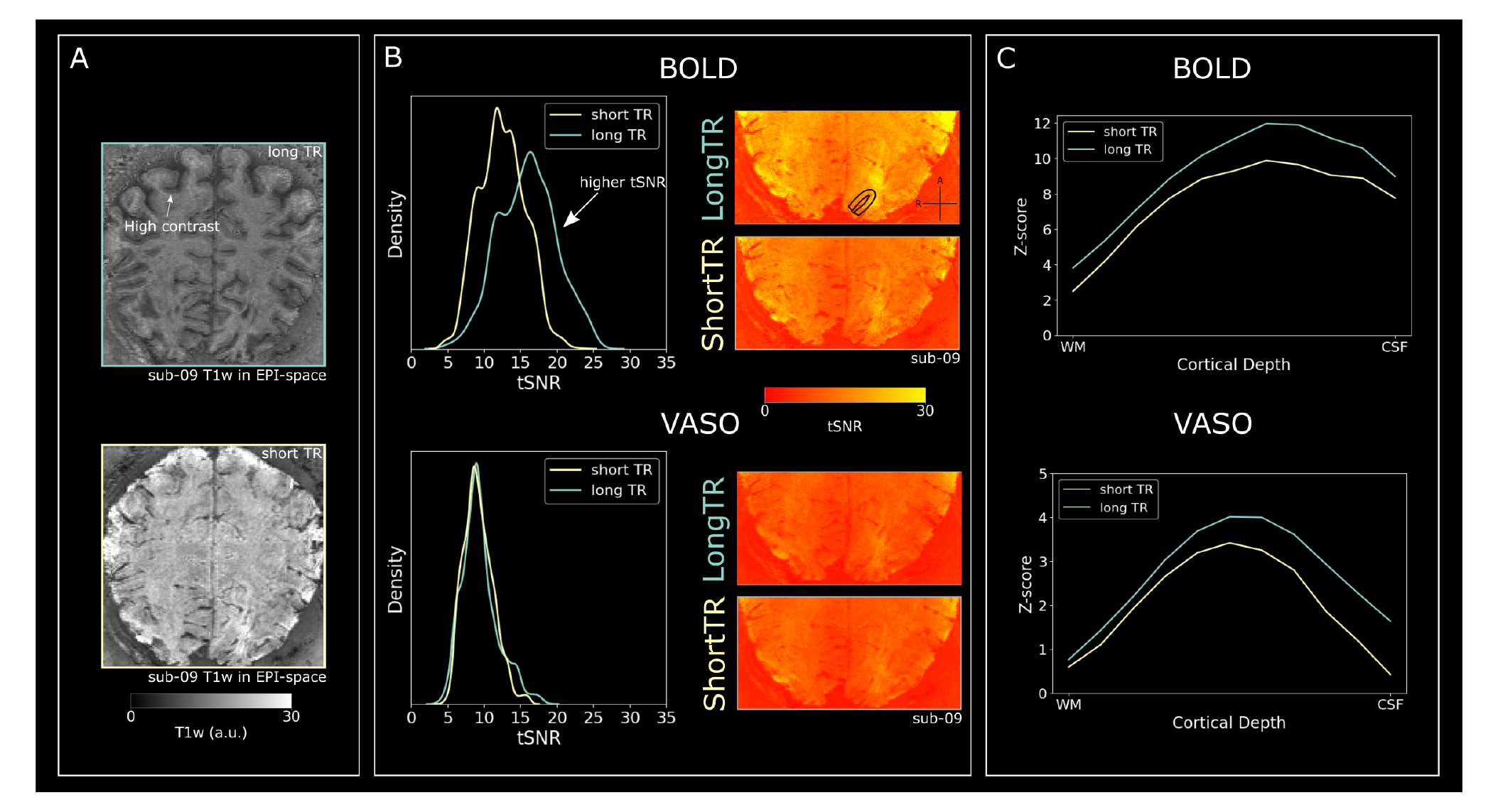
Comparison between short and long TR fMRI data in two participants. **A** T1w image in EPI-space for long (upper) and short (lower) TR fMRI data. Note the high tissue contrast between white and gray matter in the long TR data. **B** tSNR comparison between short and long TRs. Right: tSNR maps for long and short TR data. Left: Voxel-wise tSNR kernel density estimation plots (similar to smoothed histograms) for both participants together. Values were taken from V1 gray matter (example indicated in the uppermost map on the right). **C** Comparison between short and long TR data activation (z-scores, contrast: stimulation vs. rest) across cortical depths for both participants together. Values were taken from the same V1 gray matter regions as tSNR.

## Discussion

### Summary of results

In this work, we characterized the applicability of VASO for event-related paradigms. With a short 895 ms volume acquisition TR, we were able to capture the haemodynamic response for VASO and BOLD within a few (∼20) average trials. Furthermore, the short TR and high specificity of VASO enabled us to show subtle laminar-dependent timing differences of neural processing and vascular reactivity features. Specifically, superficial layers showed delayed responses in VASO but not in BOLD. Besides general advantages of event-related designs, we believe that this result demonstrates the added value of employing short stimulus paradigms using VASO, which greatly increases the range of possible research questions to be addressed with CBV-based laminar fMRI.

### Response peak location and timing across cortical depth

A distinctive feature of VASO is its microvascular weighting, which leads to strong activation of middle cortical layers during bottom-up processing (Akbari et al., 2022; Jin and Kim, 2008; Y. Yu et al., 2019). In our data, microvascular weighting of VASO seems to vary depending on stimulus duration. For block-wise stimulation we observe a peak in middle layers (**Figure 2C**). For event-wise stimulation we observe a peak between middle and superficial layers (**Figure 3C**). The subtle movement of the peak towards the superficial layers from block- to event-wise stimulation could hint at decreased microvascular weighting in VASO when employing short stimulation durations. CBV peak responses towards the superficial layers for short stimulation durations, as shown here, are in agreement with various lines of research. For example, in cats, Jin and Kim (2008) performed a time dependent analysis of responses to block-wise stimulation using GE-BOLD, CBV weighted (with monocrystalline iron oxide nanoparticle [MION] contrast agent) and CBF type imaging. They found that CBV signals are weighted towards the cortical surface at earlier stages of the response and only later develop the full weighting towards the microvasculature. Another example is Hirano et al. (2011), where they report BOLD, CBF and CBV (also measured with MION) data in rats. For stimulation durations comparable to ours, the authors find highest CBV amplitudes in superficial layers. In contrast to Hirano et al. (2011), Shen et al. (2008) report strongest CBV responses to short stimuli in middle layers. Note however, that for Shen et al. (2008) the middle layer response is only significantly different from deep but not superficial layers. Therefore, the results from Shen et al. (2008) also match our data, as we observe a peak between middle to superficial layers for event-wise stimulation.

In these preclinical studies, the relative microvascular weighting of CBV responses for short compared to long stimuli has been discussed in the context of laminar differences in the vascular architecture. Specifically, microvascular blood compartments tend to have the earliest onset times (<1 s, Silva and Koretsky, 2002; Tian et al., 2010; X. Yu et al., 2014), but take long to peak (>15 s) (Mandeville, 2012). On the other hand, larger, actively muscle-controlled arterial compartments closer to the cortical surface, tend to have later onset times (>1 s, Tian et al., 2010) but shorter times to peaks (Kennerley et al., 2012). For similar investigations of response times across CBV compartments in humans, time-resolved CBV measurements are necessary, but were not available so far.

Our event-related results indicate, for the first time, similar reactivity of CBV-compartments in humans. Specifically, we report earliest responses in deep and middle layers, paralleling the early responses in microvascular compartments found in rodent models (X. Yu et al., 2014). Furthermore, we find a delayed response-peak in superficial layers for VASO, supporting later macrovascular responses (Kennerley et al., 2012; Tian et al., 2010). To our knowledge, this is the first demonstration of laminar response time differences in humans using non-invasive CBV measurements. On the other hand, we do not find the longer time to peak for superficial layers in the BOLD data, as reported by Siero and colleagues (Siero et al., 2011; Siero et al., 2013). However, this can likely be explained by two factors. Firstly, Siero et al. report a smaller effect size. In their data, the difference in the time to peak is ∼0.23 s/mm (Siero et al., 2011). In the visual cortex, this would result in peak time differences between deep and superficial layers of ∼0.6 s. This effect is much smaller than in our VASO data (peak time differences of about 1.3 s). Secondly, the higher temporal resolution and shorter stimulus durations in their studies compared to ours. Here, we invested image encoding time in higher spatial resolutions and used longer stimuli. This allowed us to differentiate layer effects with less partial voluming of the temporal evolution of pial vessels. The subtle depth-dependent timing of BOLD- and CBV- haemodynamic responses across stimulus durations in humans could be investigated further by using systematic jitters to increase effective sampling rate.

### Considerations for fast event-related VASO

In VASO protocols with longer acquisition times (e.g. >1.5 s volume acquisition time), the inherent T1 contrast is sufficient to delineate GM/WM and GM/CSF borders in EPI space (L. Huber et al., 2015). However, in VASO protocols with shorter acquisition times (e.g. <1 s volume acquisition time), such as the one used here, the T1 contrast of the EPI images is weaker due to reduced relaxation times (**Figure 6A**). Thus, acquisition and registration of anatomical reference images is necessary. Another option for future studies might be to acquire additional VASO run(s) with longer acquisition times. For example, the acquisition of run(s) with a long TR could be straightforwardly implemented in the paradigm when acquiring independent localizers with strong tasks using block-wise stimulation. Furthermore, we saw slightly lower tSNR values in the BOLD data for short TR acquisitions (**Figure 6B**). This also translated into slightly decreased activation scores (**Figure 6C**). We estimated the decrease in detection sensitivity when employing event-related paradigms with 2 s stimulation as opposed to 30 s on/off block-designs and found a 36 and 22 % decrease in detection sensitivity for BOLD and VASO, respectively. Therefore, if event-related paradigms are crucial, and detection sensitivity is required to remain comparable to block-wise stimulation, up to 2 times longer experiments may become necessary in future event-related layer-fMRI VASO studies.

### Limitations & Future directions

We see the present study as a proof of concept that event-related stimulation is feasible using VASO. As a result, there are several aspects that can be improved in future, more extensive investigations.

In the design of our stimulation protocol, we opted for efficiency. Therefore, the inter-trial intervals we chose are rather short (3-10 s), which lead to highly overlapping haemodynamic responses between events. While this might be appropriate for neuroscience experiments in order to investigate depth-dependent differences between haemodynamic responses to different stimuli (Dale and Buckner, 1997; Glover, 1999), investigations of the haemodynamic response per se might be compromised. Rather, longer inter-trial intervals could be chosen to let the response return to baseline (van Dijk et al., 2021). This would allow researchers to account for higher-order non-linear HRF effects that are not commonly considered in the linear superposition assumption of GLM/ FIR analysis models. However, this would render the paradigm less efficient. Another aspect that is compromised by fast acquisition times at high resolutions is brain coverage. The coverage of our high temporo-spatial VASO protocol (133 × 129.542 × 114.08 mm) was sufficient to image V1 in all, and V1 and S1 in some participants. This is common for most layer-fMRI applications, which focus on single areas of interest (Schluppeck et al., 2018). However, if the goal is to study brain-wide distributed networks, this is not sufficient. Therefore, sacrificing imaging speed might be necessary to investigate connectivity between distant brain regions across cortical depth (Koiso et al., 2022).

In general, we see various routes for future applications. The most immediate benefit would be for neuroscientific applications that necessitate short TRs for efficient sampling. For example, Gau et al. (2020) report multimodal influences on the response magnitude in deep layers of the primary auditory cortex. Here, we also found indications of multimodal influences on the response timing in deep layers of V1. (**Supplementary Figure S3**). Future studies could leverage short acquisition times to investigate how these different aspects of multimodal interactions can be integrated. Furthermore, paradigms that rely on irregular perceptual switches (e.g. Schneider et al., 2019) would benefit from shorter TRs, as scan time would be dramatically decreased while still efficiently sampling different cognitive states. Finally, one of the promises of layer-fMRI is to investigate the directionality of input and output to cortical areas (L. Huber et al., 2017). Methods like Granger causality (Goebel et al., 2003) or dynamic causal modeling (Friston, 2012) may further corroborate investigations of the directional connectivity, however, these methods are highly reliant on fast sampling of responses which, to date, was not available with VASO. With our advances in short TRs in VASO, we are approaching the feasibility of these methods in future applications.

Finally, measuring different aspects of the haemodynamic response (CBF, CBV, CMRO2 and BOLD) is pivotal for its complete characterization (Uludăg et al., 2009). Siero and colleagues have investigated the haemodynamic response to short stimuli across cortical depths in humans using BOLD fMRI (Siero et al., 2015; Siero et al., 2011; Siero et al., 2013). However, the invasive nature, constrained sampling efficiency, and low SNR of CBF and CBV measurements have so far hindered thorough investigations of the CBF- and CBV- haemodynamic response at the laminar level in humans. Here, we demonstrate laminar response time differences in humans using non-invasive CBV measurements. We believe that with our implementation of fast VASO acquisition, we have enabled crucial investigations of the cortical depth-dependent evolution of biophysical hemodynamic (Havlicek and Uludăg, 2020; Puckett et al., 2016) and neurophysiological processes (Petridou and Siero, 2019).

## Conclusion

High-resolution VASO has proven to be a robust method to study brain function across cortical depth. For an even wider application in neuroscientific research, fast acquisition schemes and stimulation protocols are crucial. Huettel (2012) commented on the impact of event-related designs as follows: “No other advance—not stronger magnetic fields, not improved pulse sequences, nor even sophisticated new analyses—has contributed more to the popularization of fMRI than event-related approaches to experimental design” (Huettel, 2012). Indeed, layer-fMRI VASO is highly dependent on strong magnetic fields, improved pulse sequences and sophisticated analyses. Now, with this study, we hope to have contributed to the further popularization of VASO by showing that fast event-related designs are possible and provide meaningful insights.

## Acknowledgments

Data was acquired at Scannexus (Maastricht, the Netherlands). We thank Chris Wiggins for providing the 3rd order shimming tools used here. We thank Federico DeMartino for discussions on the analysis of event-related fMRI data. We thank Vojťech Smekal for helpful discussions on the manuscript. Special thanks goes to the remaining “Maastricht layer-seminar” members Lonike Faes, Miriam Heynckes, Kenshu Koiso, Alessandra Pizzuti and Yawen Wang for countless discussions on topics related to layer-fMRI. We thank Benedikt Poser for kindly sharing his 3D-EPI sequence code used here and helpful advice on layer-fMRI VASO optimization and reconstructions. Finally, we thank Dr. Amanda Kaas, who was supervising Sebastian Dresbach in the beginning of this project and provided the piezo-electric stimulation setup.

Sebastian Dresbach is supported by the ‘Robin Hood’ fund of the Faculty of Psychology and Neuroscience and the department of Cognitive Neuroscience. Laurentius Huber was funded by the NWO VENI project016.Veni.198.03 and the York-Maastricht Partnership. Scan time was kindly provided by Maastricht University Faculty of Psychology and Neuroscience via the intramural MBIC funding scheme. Rainer Goebel is partly funded by the European Research Council Grant ERC-2010-AdG269853 and Human Brain Project Grant FP7-ICT-2013-FET-F/604102. We thank Brain Innovation for supporting this project and for funding Omer Faruk Gulban while working on it.

## Data and Software availability statement

Analysis code is available on github: <https://github.com/sdres/eventRelatedVASO>. Data is available on OpenNeuro: <https://openneuro.org/datasets/ds004539>.

## Author Contributions

According to the CRediT (Contributor Roles Taxonomy) system.

|  |  |
| --- | --- |
| Conceptualization | S.D., R.H., R.G. |
| Data curation | S.D., R.H. |
| Formal analysis | S.D. |
| Funding acquisition | R.G. |
| Investigation | S.D. R.H. R.G. |
| Methodology | S.D. R.H. O.F.G. R.G. |
| Project administration | S.D. R.H. R.G. |
| Resources | S.D. R.H. R.G. |
| Software | S.D. R.H. O.F.G. R.G. |
| Supervision | R.H. R.G. |
| Validation | S.D. |
| Visualization | S.D. R.H. O.F.G. R.G. |
| Writing - original draft | S.D. |
| Writing - review and editing | S.D. R.H. O.F.G. R.G. |

## Supplementary figures

**Figure S1:**
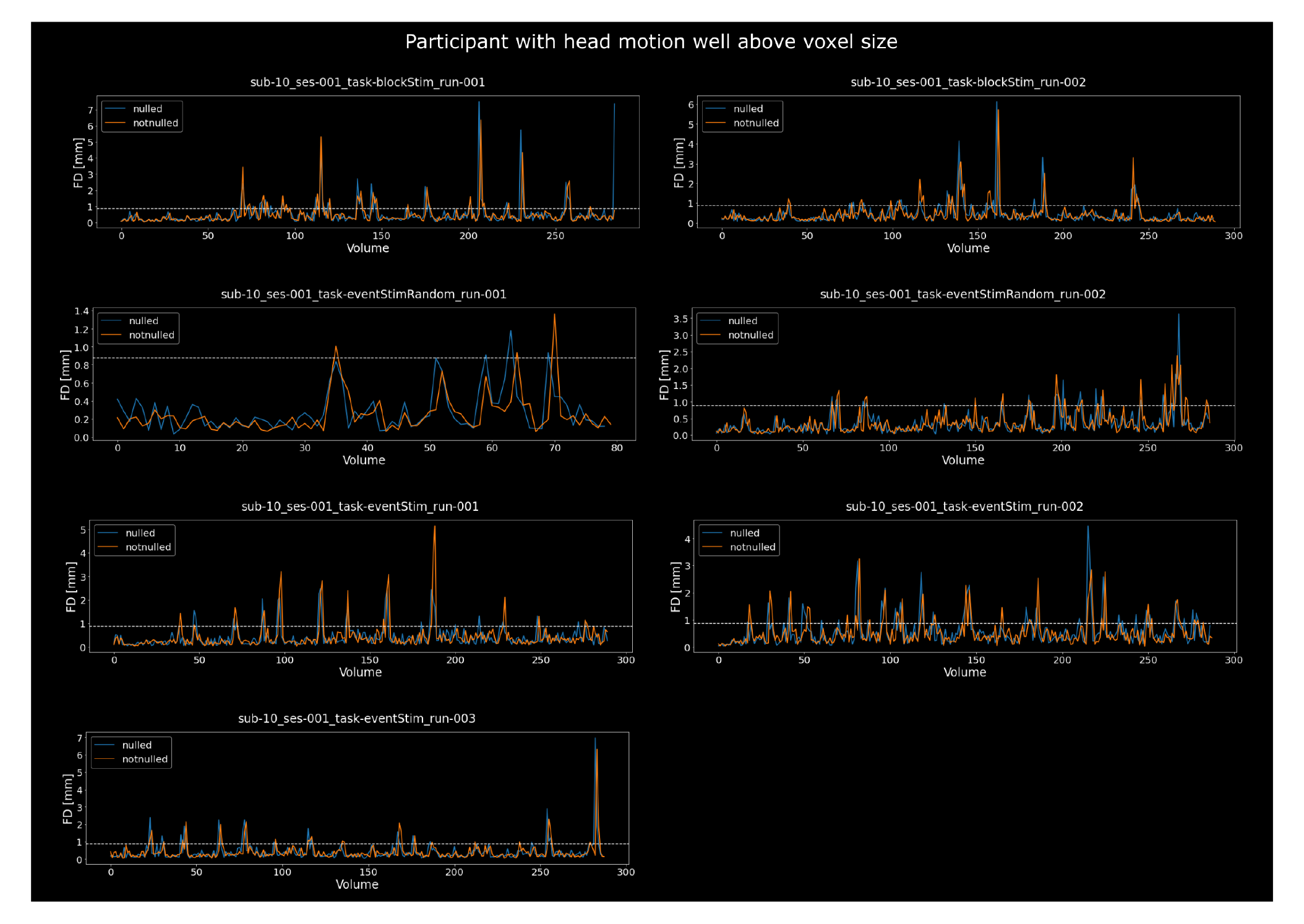
Participant with excessive head motion. Framewise displacements (FDs) for each run of the only participant consistently showing motion peaks greater than a voxel size (0.88mm, indicated by white dashed line). This participant was therefore excluded from further analyses.

**Figure S2:**
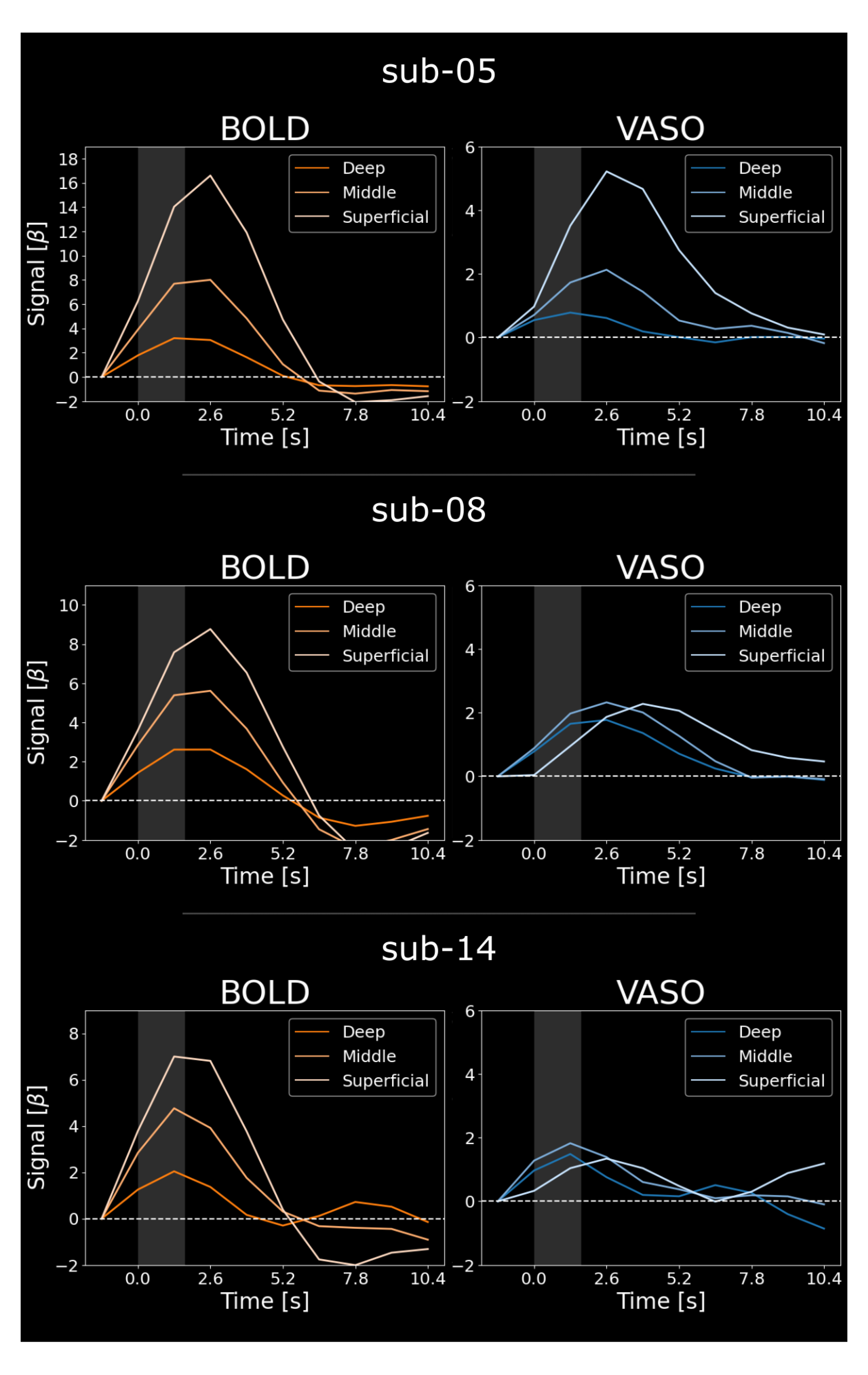
FIR-model results show high inter-participant variability with respect to peak layer. Same as **Figure 3B** but for 3 individual participants (upper: sub-05, middle: sub-08, lower: sub-14). Note that the superficial layer VASO activity varies across participants. While sub-08 resembles the group results, sub-05 shows greatest activity in superficial layers and sub-14 shows lowest responses in superficial layers. Nonetheless, the delayed peak for superficial layers is preserved in all participants.

**Figure S3:**
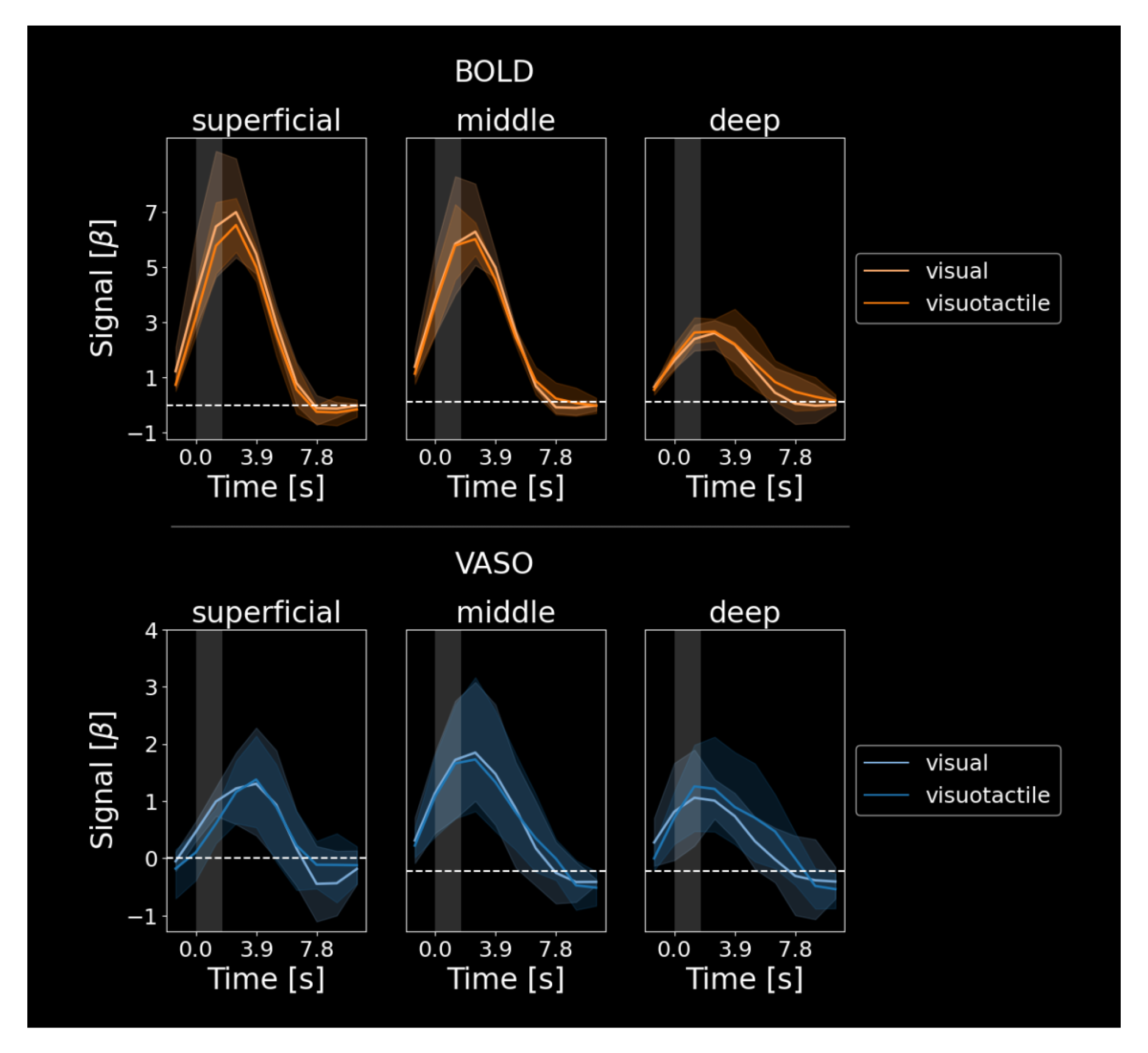
Comparison between visual and visuo-tactile stimulation in V1. BOLD (upper) and VASO (lower) activation (n = 4) in response to visual and visuo-tactile stimulation separately. Data is plotted separately for the three layers (deep, middle, superficial). Visual and visuo-tactile stimulation evoke very similar responses in superficial and middle layers for BOLD and VASO. In deep layers, visuo-tactile stimulation shows a slightly prolonged response compared to visual stimulation only. This effect was present in the BOLD and VASO data, while being more pronounced in the latter. However, this effect is rather small (within error bars) and has to be interpreted with caution. Still, we believe that this might indicate a multisensory integration effect taking place in deep layers of V1 in response to additional tactile input. Future studies on neuroscientific event-related applications might investigate this further.

